# Increases in structural shortest path lengths provide information about the distal neurophysiological consequences of focal brain lesions

**DOI:** 10.1101/785576

**Authors:** Joseph C. Griffis, Nicholas V. Metcalf, Maurizio Corbetta, Gordon L. Shulman

## Abstract

Focal brain lesions disrupt resting-state functional connectivity, but the underlying structural mechanisms are unclear. Here, we examined the direct and indirect effects of structural disconnections on resting-state functional connectivity in a large sample of sub-acute stroke patients with heterogeneous brain lesions. We defined direct disconnections as the loss of direct structural connections between two regions, and indirect disconnections as increases in the shortest structural path length between two regions that lack direct structural connections. On average, nearly 20% of all region pairs suffered either a direct or indirect disconnection by the lesions in our sample. Importantly, both directly and indirectly disconnected region pairs showed more severe functional connectivity disruptions than region pairs with spared direct and indirect connections, respectively, although functional connectivity disruptions tended to be most severe between region pairs that sustained direct structural disconnections. Together, these results emphasize the widespread impacts of focal brain lesions on the structural connectome, and show that these impacts are reflected by disruptions of the functional connectome. Further, they indicate that in addition to direct structural disconnections, lesion-induced increases in the structural shortest path lengths between indirectly structurally connected region pairs provide information about the remote functional disruptions caused by focal brain lesions.

## 1. Introduction

Focal brain lesions that result from stroke and other neurological disorders produce widespread disruptions of brain function that often involve regions remote from the site of injury (Carrera and Tononi, 2014). The distributed functional consequences of focal brain lesions can be measured non-invasively using resting-state fMRI functional connectivity, a measure of the correlation between ongoing low-frequency blood oxygen-level dependent signal fluctuations in different brain regions (Biswal et al., 1995). Lesion-induced functional connectivity disruptions have been strongly associated with clinical deficits in multiple cognitive and behavioral domains (Baldassarre et al., 2016), but it remains unclear how they relate to the structural impact of the underlying lesion. Clarifying the nature of this relationship is thus an important goal that has the potential to inform the development of new therapeutic approaches that aim to restore normal brain function using techniques such as non-invasive neurostimulation (Carrera and Tononi, 2014; Raffin and Siebner, 2014), and beyond clinical relevance, also represents an important step towards understanding how brain structure shapes brain function.

We recently made progress towards this goal by demonstrating that the severity of commonly observed network-level functional connectivity disruptions observed after stroke, namely reductions of interhemispheric integration within networks and ipsilesional segregation between networks, are more strongly related to lesion-induced structural disconnections of inter-regional white matter pathways than to lesion size, location, or damage to putative critical grey matter regions (Griffis et al., 2019). However, our results only hinted at the specific structural features that provide information about when functional connectivity disruptions are sustained between region pairs. For example, we observed significant, albeit weak-to-moderate, connection-level correspondences between the structural disconnection and functional connectivity patterns that co-varied together across patients, suggesting that a lesion that interrupts the direct structural connection between two regions also tends to disrupt their functional connectivity. Here, we explicitly tested whether region pairs that suffer a direct structural disconnection reliably show more severe disruptions of functional connectivity than region pairs with spared direct structural connections.

Most lesion-induced disruptions of functional connectivity, however, occur between regions that can only be connected by traversing intermediate structural links. This suggests that many functional connectivity disruptions may reflect the up/downstream consequences of direct structural disconnections. Supporting this view, a previous study of eleven stroke patients reported that pontine lesions, which should damage intermediary connections along the polysynaptic cortico-pontine-cerebellar pathway, disrupted functional connectivity between somatomotor cortex and the contralateral cerebellar hemisphere (Lu et al., 2011). Therefore, we also sought to identify a structural feature that provides information about the functional connectivity disruptions sustained by indirectly structurally connected region pairs. The report by Lu et al., (2011) suggests that disruptions of the shortest structural paths between indirectly structurally connected region pairs might represent such a feature. Importantly, shortest structural path lengths (SSPLs), which correspond to the minimum number of direct structural links that must be traversed to establish a structural pathway connecting a region pair, have been previously shown to influence functional connectivity in healthy individuals (Goni et al., 2014).

Here, we tested whether damage to the intermediary structural connections along the shortest structural paths linking indirectly structurally connected region pairs represents a general mechanism of lesion-induced functional connectivity disruptions across the cortex. Specifically, we tested whether indirectly structurally connected regions that exhibit lesion-induced increases in SSPLs show disruptions of functional connectivity attributable to this putative indirect structural disconnection (see **Fig. 1**). Because a lesion that interrupts the direct structural connection between two regions also increases their SSPL (i.e. from one to greater than one), the SSPL measure allowed a unified treatment of both directly and indirectly structurally connected region pairs (**Fig. 1**). Accordingly, we also compared the magnitudes of the functional connectivity disruptions associated with direct vs. indirect structural disconnections.

**Fig. 1.**
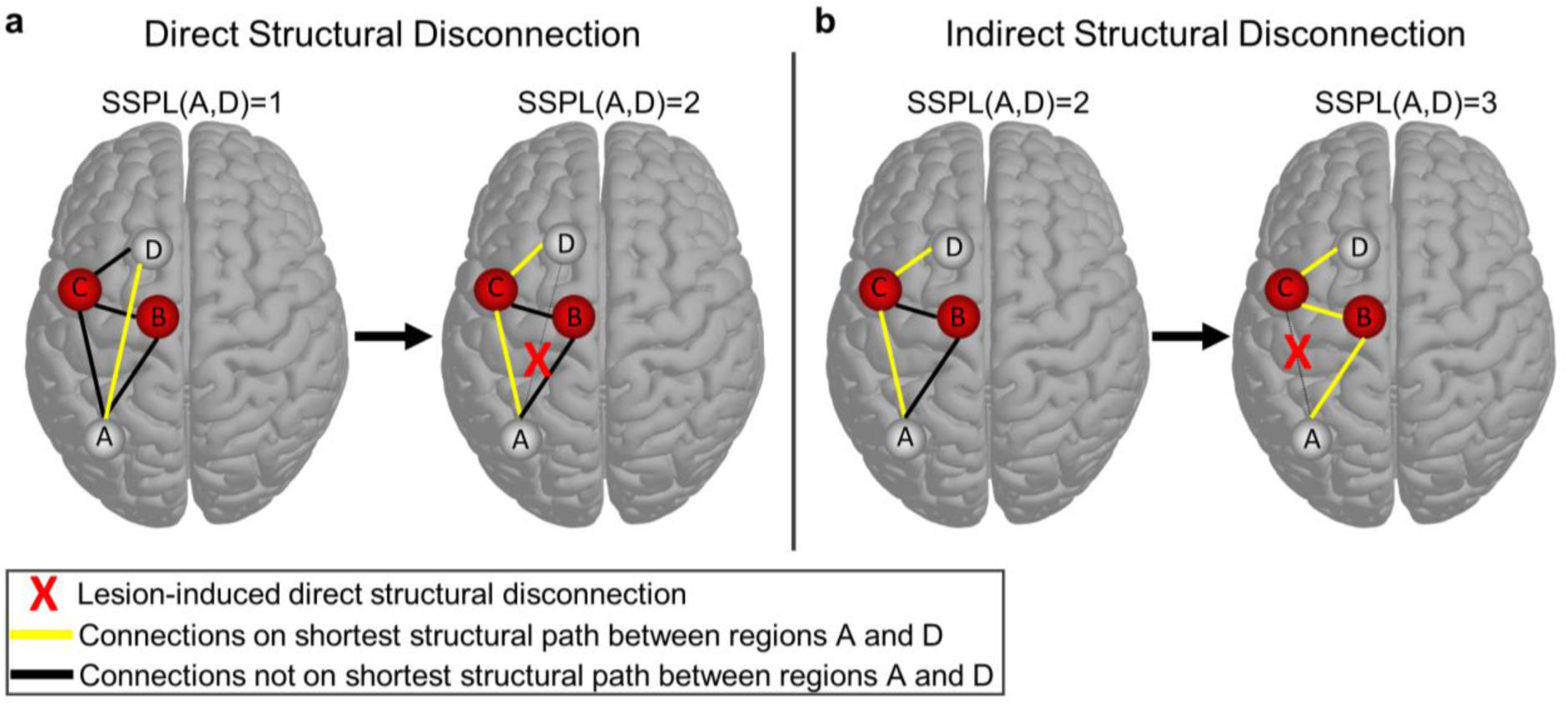
Defining direct and indirect structural disconnections. **(a)** The brain on the left shows a simple network where regions *A* and *D* are directly structurally connected to each other (yellow line), and therefore have an SSPL equal to 1. The brain on the right shows the SSPL (yellow line) between regions *A* and *D* after the direct structural connection has been disrupted by a lesion (red X): the SSPL between regions *A* and *D* is now 2 because the shortest path passes through region *C*. This is an example of how direct structural disconnections increase SSPLs between disconnected regions. **(b)** The brain on the left shows an alternative network configuration where regions *A* and *D* are indirectly structurally connected to each other via mutual connections to region *C* (yellow line), and therefore have an SSPL equal to 2. The brain on the right shows the SSPL (yellow line) between regions *A* and *D* after the structural connection between regions *A* and *C* has been disrupted by a lesion (red X): the SSPL between regions *A* and *D* is now 3 because the shortest path passes through both regions *B* and *C*. This is an example of how a direct structural disconnection can increase SSPLs between indirectly structurally connected regions, which we refer to here as an “indirect structural disconnection”.

## 2. Methods

### 2.1. Participant information

Written informed consent was obtained prior to enrolling subjects in the study. Study procedures were performed in accordance with the Declaration of Helsinki ethical principles and approved by the Institutional Review Board at Washington University in St. Louis. The complete data collection protocol is described in complete detail elsewhere (Corbetta et al., 2015). Data from 132 first-time stroke patients with clinical evidence of impairment and data from 36 demographically matched healthy controls were considered for inclusion in the analyses. After applying quality controls (described in Section 2.7), data from a total of 114 patients and 24 controls were included in the study. Demographic information is provided in **Table 1. Data and code availability statement:** The full set of neuroimaging data (along with behavioral data) are available at http://cnda.wustl.edu/app/template/Login, and specific data and scripts are available upon request to the authors.

**Table 1.** Participant Demographics.

| Group | Age (Mean/SD) | Sex | Handedness | Education (Mean/SD) | Lesion Side |
| --- | --- | --- | --- | --- | --- |
| Patients (N=114) | 52.4/10.2 | 58 F, 56 M | 106 R, 8 L | 13.3/2.8 | 54 R, 60 L |
| Controls (N=24) | 54.5/13.5 | 12 F, 12 M | 23 R, 1 L | 13.5/2.1 | N/A |

SD: Standard Deviation, M: Male, F: Female, R: Right, L: Left

### 2.2. Neuroimaging data collection

Neuroimaging data were collected at the Washington University School of Medicine using a Siemens 3T Tim-Trio scanner with a 12-channel head coil. We obtained sagittal T1-weighted MP-RAGE (TR=1950 msec; TE=2.26 msec, flip angle = 90 degrees; voxel dimensions = 1.0×1.0×1.0 mm), transverse T2-weighted turbo spin-echo (TR=2500 msec; TE=43 msec; voxel dimensions = 1×1×1), sagittal T2-weighted FLAIR (TR=750 msec; TE=32 msec; voxel dimensions = 1.5×1.5×1.5 mm) and gradient echo EPI (TR=2000 msec; TE=2 msec; 32 contiguous slices; 4x4 mm in-plane resolution) resting-state functional MRI scans from each subject. Participants were instructed to fixate on a small centrally-located white fixation cross that was presented against a black background on a screen at the back of the magnet bore. An Eyelink 1000 eye-tracking device (SR Research) was used to monitor eye status (i.e. eyes opened/closed) during each functional MRI run. We obtained between six and eight resting-state scans (128 volumes each) from each participant (∼ 30 minutes total).

### 2.3. Lesion identification

The Analyze software package (Robb and Hanson, 1991) was used to manually segment lesions on structural MRI. The different structural scans were used in conjunction to ensure that lesions were completely segmented. Surrounding vasogenic edema was included in the lesion definition if present. Two board certified neurologists (Maurizio Corbetta and Alexandre Carter) reviewed all segmentations, and segmentations were also reviewed a second time by MC. The final segmentations were converted to binary lesion masks, transformed to MNI atlas space using both linear and non-linear registrations, and were resampled to have isotropic voxel resolution.

### 2.4. Regional parcellation

We parcellated the cortex according to the Gordon333 resting-state functional MRI parcellation. This parcellation contains 333 cortical regions that are each assigned to 1 of 13 large-scale resting-state networks (Gordon et al., 2016). As in previous analyses of this dataset, 9 regions were excluded for having very low numbers of vertices (Griffis et al., 2019; Siegel et al., 2018, 2016). Surface-based functional connectivity analyses were performed using the remaining 324 regional parcels.

A set of 35 sub-cortical and cerebellar regions were also defined to enable a thorough quantification of damage and disconnection. 34 regions corresponding to portions of the cerebellum, thalamus, and basal ganglia were taken from the automatic anatomical labeling (AAL) atlas (Tzourio-Mazoyer et al., 2002). An additional region corresponding to the brainstem was taken from the Harvard-Oxford Subcortical Atlas. The “dilate” command in DSI_studio was used to dilate the cortical regions by 2mm to increase the sensitivity of subsequent region-to-region structural connectivity analyses (Van Den Heuvel et al., 2009; Wilson et al., 2011). This also led to a more liberal definition of cortical damage, since the dilated regions extended slightly beneath the pial surface (Pustina et al., 2017). The volume-based regional parcels are illustrated in **Figure 2b** (left), and the surface-based cortical parcels are illustrated in **Figure 3a**.

**Fig. 2.**
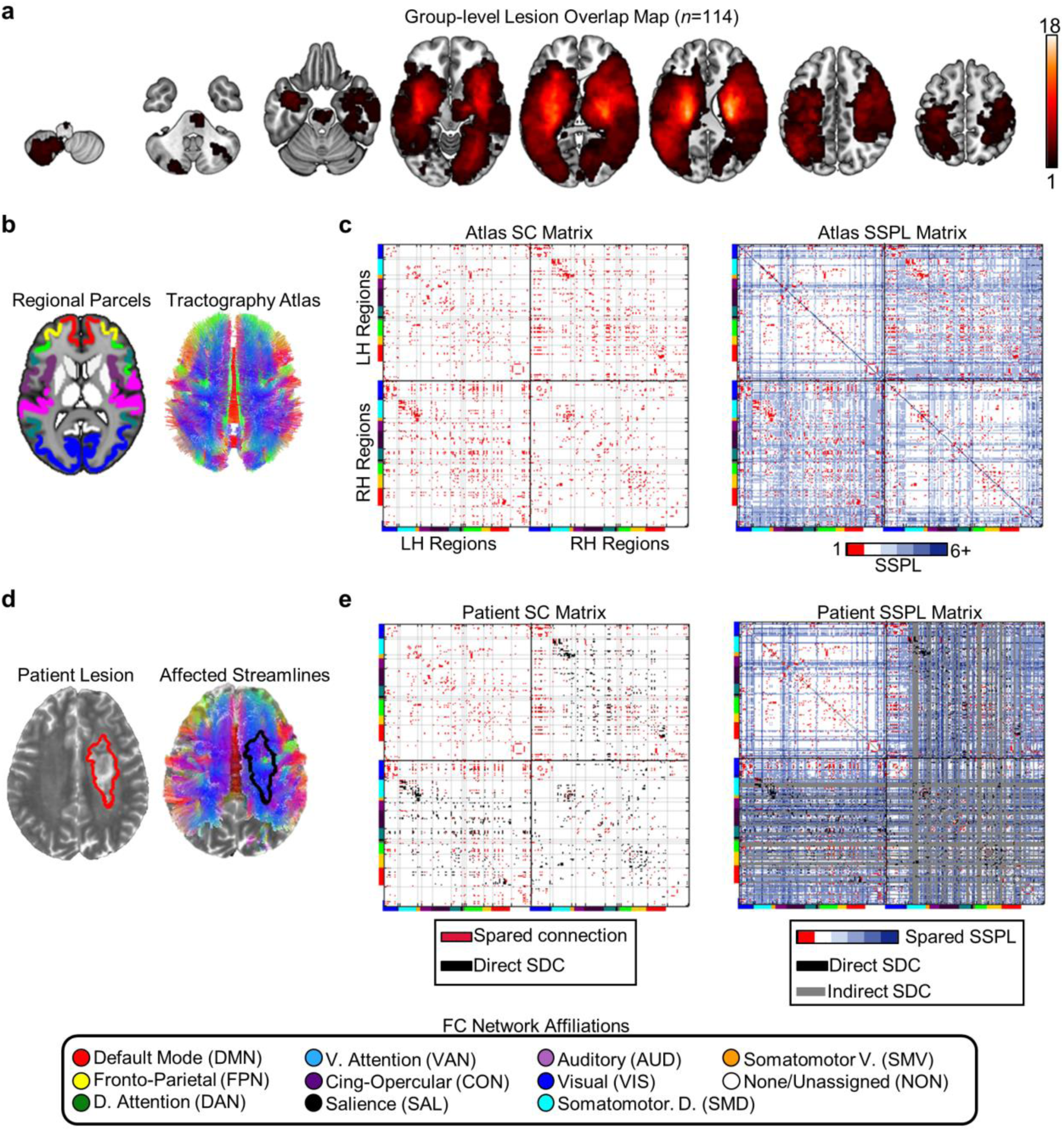
Structural data. **(a)** The colormap shows the number of patients with overlapping lesions at each voxel in the brain. The color scale ranges from the minimum (i.e. 1) to the maximum (i.e. 18) lesion overlap across patients (*n*=114). **(b)** The regional parcels and tractography atlas. **(c)** Left – Cortical SC matrix derived from the tractography atlas and regional parcels. Right – Cortical SSPL matrix derived from the atlas SC matrix. Directly structurally connected regions have SSPLs equal to 1. **(d)** Lesion segmentation (left) and affected streamlines (right) for a single patient. **(e)** Left --spared cortical direct structural connections (red) and direct cortical structural disconnections (black – direct SDC) for the patient shown in (d). Right – spared cortical SSPLs (colored), indirect cortical structural disconnections (gray – indirect SDC), and direct cortical structural disconnections (black– direct SDC) for the same patient shown in (d). Note that the upper left quadrants in (d) and (e) correspond to contralesional intrahemispheric connections, which were spared by the right hemisphere lesion. The colored bars on matrix axes correspond to the cortical functional connectivity network assignments of each region shown in the legend at the bottom of the figure. Note: SSPL calculations also considered subcortical/cerebellar connections (not shown).

**Fig. 3.**
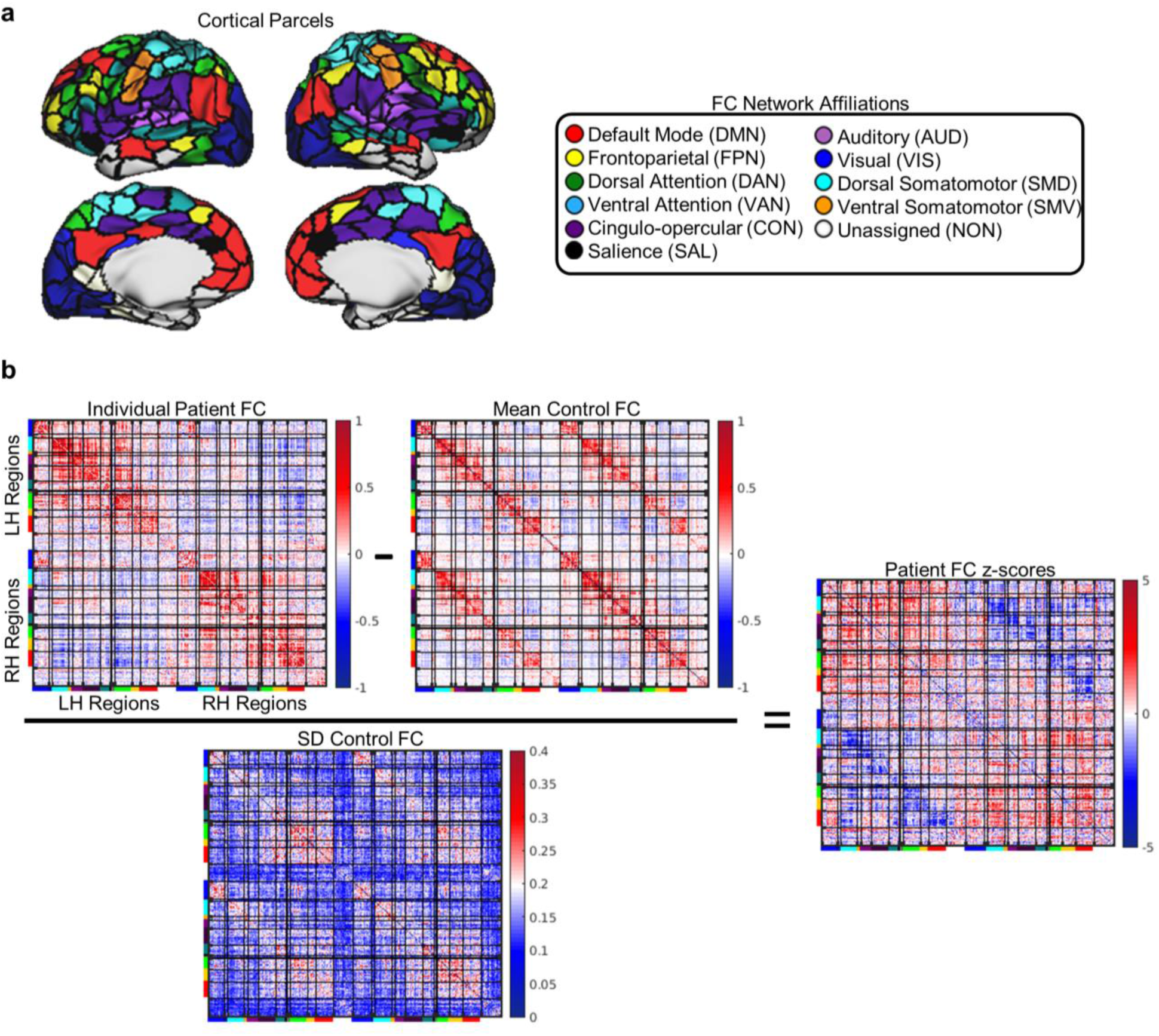
Functional connectivity disruptions. **(a)** Cortical regions are shown on the cortical surface and are color-coded by network assignment. **(b)** The transformation of a raw functional connectivity matrix to a functional connectivity *z*-score matrix is illustrated for the same patient shown in **Fig. 2d-e**. The bars on matrix axes correspond to the cortical functional connectivity network assignments shown in the legend at the top of the figure.

### 2.5 Structural connectome atlas

We created a structural connectome atlas using a publicly available diffusion MRI streamline tractography atlas based on high angular resolution diffusion MRI data collected from 842 healthy Human Connectome Project participants (Yeh et al., 2018) (**Fig. 2b**, right). These data were accessed under the WU-Minn HCP open access data use term. The tractography atlas contained expert-vetted streamline trajectories in MNI space, and each streamline was assigned to 1 of 66 macroscale white matter pathways (e.g. arcuate fasciculus, cortico-spinal tract, corpus callosum, etc.).

MATLAB scripts implementing functions from the DSI_studio software package was used to define the normative pairwise structural connectome based on the tractography atlas (**Fig. 2c**, left). We first combined the various streamline bundles from the tractography atlas (e.g. short-range U-fibers, callosal projections, etc.) into a single. *trk* file that included all streamlines associated with each of the fiber pathways in the atlas. We then extracted all pairwise structural connections, defined as streamlines that bilaterally terminated (i.e. began and ended) within any pair of the 359 volume-based regions, to create a 359×359 structural connectivity adjacency matrix where each cell quantified the number of streamlines connecting a region pair. The close proximity of ventral visual and dorsal cerebellar regions led to a few dorsal cerebellar streamlines terminating within the dilated ventral visual regions, and so any connections between the visual cortex and cerebellum were removed. A breadth-first search algorithm, implemented in the *breadthdist* function provided with the Brain Connectivity Toolbox (Rubinov and Sporns, 2010) for MATLAB was then applied to the binarized structural connectivity matrix to obtain a matrix of pairwise SSPLs (**Fig. 2c**, right), where each cell quantified the minimum number of structural connections that needed to be traversed to establish a structural path between each region pair (Goni et al., 2014). A total of 15 cortical regions had undefined SSPLs, and so they were excluded from further analyses.

### 2.6. Structural connectivity measures

MATLAB scripts implementing functions from the DSI_studio software package were used to estimate the expected direct structural disconnections for each patient by intersecting the MNI-registered lesion mask with streamline tractography atlas (**Fig. 2d**). For each patient, all streamlines that intersected the lesion were extracted into a 359×359 direct structural disconnection matrix where each cell quantified the number of streamlines between each region pair that intersected the lesion (**Fig. 2e**). The direct structural disconnection matrix was then subtracted from the atlas-derived structural connectivity matrix to obtain a spared structural connection matrix where each cell quantified the number of streamlines between each region pair that were spared by the lesion. The structural disconnection and spared structural connection matrices were then normalized by dividing each cell by the corresponding number of streamlines in the atlas structural connectivity matrix so that each cell in the normalized spared structural connection and direct structural disconnection matrices quantified the proportion of streamlines between each region pair that was spared vs. directly disconnected by the lesion, respectively. Each matrix was then binarized by setting all non-zero values to 1.

For each patient, a patient-specific SSPL matrix was created by applying the *breadthdist* function to that patient’s spared connection matrix (**Fig. 2e**). The atlas SSPL matrix was then subtracted from the patient-specific SSPL matrix to obtain an SSPL difference matrix. The SSPL difference matrix was then thresholded and binarized so that cells corresponding to region pairs with preserved SSPLs had values equal to 0 and cells corresponding to region pairs with increased SSPLs had values equal to 1. Values for all cells that corresponded to structurally connected region pairs were set to 0 so that 1-valued cells in the resulting matrix corresponded to structurally un-connected region pairs that sustained indirect disconnections. A spared indirect connection matrix was then created where all cells that corresponded to structurally un-connected region pairs with spared SSPLs had values equal to 1. All subsequent analyses involving these matrices utilized only the above-diagonal elements (i.e. upper triangles) of each matrix.

### 2.7. fMRI data processing

The following steps were used to preprocess the fMRI data: slice-timing correction using sinc interpolation, correction of inter-slice intensity differences resulting from interleaved acquisition, normalization of whole-brain intensity values to a mode of 1000, correction for distortion via synthetic field map estimation, and within- and between-scan spatial re-alignment. The fMRI data were re-aligned, co-registered to the corresponding structural scans, registered to atlas space using both linear and nonlinear registration procedures, and resampled to 3mm cubic voxel resolution.

Additional preprocessing was employed to remove contributions from non-neural sources of signal variance. The six head motion parameters obtained from rigid body correction were regressed from the data along with the global whole-brain signal and signals from CSF and white matter tissue compartments that were extracted using FreeSurfer tissue segmentations (Dale et al., 1999; Fischl et al., 1999). Band-pass filtering (0.009 < *f* < 0.08 Hz) was applied to retain low-frequency fluctuations. Frame censoring was applied using a 0.5mm framewise displacement threshold, and frames that succeeded high-motion frames were also censored to reduce artifacts related to head motion (Power et al., 2014). For each run, the first four frames were discarded to allow for the scanner to reach steady-state magnetization.

Cortical surfaces were generated and fMRI data were further processed according to previously published procedures (Glasser et al., 2013) with some modifications to improve processing of lesioned brains (Siegel et al., 2017). Anatomical surfaces were obtained from the T1-weighted scans using Freesurfer (Dale et al., 1999; Fischl et al., 1999) and were visually inspected to ensure quality. For patients with failed registrations/segmentations, the T1-weighted scans were modified by replacing lesioned voxels with normal tissue voxels from the structural atlas. The registration/segmentation procedures were then re-run, and the modified voxels were masked out after running the procedures (Siegel et al., 2017).

Each hemisphere was resampled to 164,000 vertices, and the two hemispheres were registered to each other (Van Essen et al., 2001) before down-sampling to 32,000 vertices per hemisphere. Ribbon-constrained sampling (implemented in Connectome Workbench) was utilized to sample each subject’s fMRI data on the corresponding individual surface. Any voxels with coefficients of variation > 0.5 standard deviations above the mean within a 5 mm sigma Gaussian neighborhood were not included in the mapping from volume to surface (Glasser et al., 2013).

As in our previous work (Griffis et al., 2019), subjects were included if they had at least 180 usable frames of functional MRI data. Data from a total of 18 patients and 9 controls were excluded, and data from the remaining 114 patients and 24 controls were included in subsequent analyses.

### 2.8. Functional connectivity

Pairwise functional connectivity matrices were constructed by correlating the mean (i.e. across all within-region vertices) post-processed BOLD timeseries from each surface region with that of each other region in the brain. Fisher z-transformation was then applied to the resulting linear correlation values to obtain 324×324 pairwise functional connectivity matrices for each patient and control. Any vertices that fell within the boundaries of the lesion were excluded from functional connectivity estimation. Any regions with less than 60 vertices remaining after accounting for the lesion were completely excluded by setting the corresponding values to NaN as in previous work (Griffis et al., 2019; Siegel et al., 2018, 2016). All subsequent analyses were performed using only the above-diagonal elements (i.e. upper triangle) of the functional connectivity matrices.

Patient functional connectivity matrices were converted to z-score matrices by subtracting the control mean functional connectivity matrix and dividing the result by the control standard deviation functional connectivity matrix (**Fig. 3b**). Each cell in the resulting matrices therefore quantified the distances (in standard deviation units) between the observed patient functional connectivity values and the expected values from the control group.

### 2.9. Statistical Analyses

Prior to performing analyses using the patient data, we aimed to replicate the previously reported dependence of normal functional connectivity on SSPLs using data from the control group (**Fig. S2**). Each functional connection in the brain was categorized into one of four categories based on its network (i.e. within-network vs. between-network) and hemispheric (i.e. interhemispheric vs. intrahemispheric) connection type. For each control participant, we extracted the mean functional connectivity values for region pairs with SSPLs equal to 1, 2, 3, and 4+ within each connection type category. For each connection type category, we then performed one-way repeated measures ANOVAs with a factor of SSPL to determine whether mean functional connectivity for that connection type category varied systematically across SSPLs in the control group (**Fig S2a**). Greenhouse-Geisser correction was applied to the degrees of freedom to account for violations of the sphericity assumption (Abdi, 2010). These analyses were also repeated after regressing out the effects of log-transformed Euclidean distances on functional connectivity (**Fig. S2b**). The log transformation was applied because this increased the linearity of the relationship between Euclidean distances and mean functional connectivity in the control group (transformed: *r*=-0.45, untransformed: *r*=-0.37).

For each patient, the mean functional connectivity z-scores were separately computed for sets of regions with spared direct structural connections, direct structural disconnections, spared indirect connections, and indirect structural connections with positive vs. negative signs in the control mean functional connectivity matrix (see **Fig. 2**). The resulting values were then entered into a three-way repeated measures ANOVA with factors of normative functional connectivity sign, connection type, and connection status. Greenhouse-Geisser correction was again applied to the degrees of freedom to account for violations of the sphericity assumption. Dependent samples *t*-tests were used to follow up the significant three-way interaction effect. Bonferroni-Holm correction was used to control the family-wise error rate at 0.05 (Aickin and Gensler, 1996). These analyses were also performed after summarizing patient-level functional connectivity z-scores by different network (i.e. within-network and between-network) and hemispheric (i.e. interhemispheric and intrahemispheric) connection types (**Fig. S3**).

Additional analyses were performed to determine the influence of arbitrary parameter choices on our results (**Fig. S4**). To ensure that our results were not dependent on the inclusion of partially damaged parcels in the main analyses, we repeated the analyses while excluding all damaged parcels (**Fig. S4a**). To ensure that our results were not dependent on the threshold for determining direct structural disconnections, we repeated the analyses using a 50% threshold for determining direct structural disconnections (**Fig. S4b**). To ensure that our results were not dependent on the inclusion of patients with severe hemodynamic lags, we repeated our analyses after excluding patients with mean interhemispheric hemodynamic lag differences greater than 2 standard deviations from the control mean (**Fig. S4c**). Finally, to ensure that our results were not dependent on the use of healthy controls as the reference group for computing functional connectivity *z*-scores (i.e., that they did not reflect a general consequence of stroke), we repeated the analyses using a subset of 22 patients who did not sustain any direct cortico-cortical disconnections as the reference group (**Fig. S4d**). These additional analyses all produced highly similar results to the main analyses reported in the text, indicating that our results did not depend on these analysis choices.

## 3. Results

### 3.1. Measuring the direct and indirect disconnections caused by stroke

We used structural and functional MRI data from a sample of 132 sub-acute (mean time post-stroke=13.57 days, SD=4.95 days) stroke patients and 36 demographically matched healthy controls that were collected as part of a larger study on stroke recovery (Corbetta et al., 2015) to study the effects of direct and indirect structural disconnections on resting-state functional connectivity after stroke. Data from 114 patients and 24 controls met our quality control criteria and were included in the study. Participant demographics are provided in **Table 1**, and the group-level lesion topography (*n*=114) is summarized in **Figure 2a**.

As in our prior work, we used an atlas-based approach to estimate the direct structural disconnections caused by each patient’s lesion (Griffis et al., 2019). We first constructed a template structural connectome by dividing the brain into 359 regional parcels (Gordon et al., 2016; Tzourio-Mazoyer et al., 2002) (**Fig. 2b**, left) and extracting all pairwise direct structural connections from a diffusion MRI streamline tractography atlas that was constructed using data from 842 healthy Human Connectome Project (HCP) participants (Yeh et al., 2018) (**Fig. 2b**, right). We then binarized the resulting direct structural connectivity matrix so that it encoded the presence vs. absence of direct structural connections between each region pair (**Fig. 2c**, left). A breadth-first search algorithm (Rubinov and Sporns, 2010) was then applied to the binarized direct structural connectivity matrix to derive a matrix of normative pairwise SSPLs (**Fig. 2c**, right). The resulting direct structural connectivity and SSPL matrices therefore represented the normative layout of inter-regional connections in the structural connectome.

We estimated the direct structural disconnections caused by each patient’s lesion (**Fig. 2d**, left) by extracting all pairwise atlas direct structural connections that intersected the lesion volume (**Fig. 2d**, right), resulting in a matrix of pairwise direct structural disconnections (**Fig. 2e**, black cells in left/right matrices). Any pairwise direct structural connections that did not intersect the lesion volume were considered to be spared by the lesion (**Fig. 2e**, red cells in left matrix) as illustrated in **Figure 1a**. To estimate the indirect structural disconnections caused by each patient’s lesion, the breadth-first search algorithm was applied to the spared direct connection matrix to derive a matrix of pairwise SSPLs (**Fig. 2e**, right matrix). We then subtracted the resulting patient-specific SSPL matrices from the atlas-derived SSPL matrix, and defined indirectly structurally connected region pairs that showed increased SSPLs relative to the corresponding atlas SSPLs as indirectly structurally disconnected (**Fig. 2e**, grey cells in right matrix). Any SSPLs between indirectly structurally connected region pairs that remained unchanged relative to the atlas SSPL matrix were considered to be spared by the lesion (**Fig. 2e**, colored cells in right matrix) as illustrated in **Figure 1b**.

### 3.2. Lesions have widespread impacts on direct and indirect structural connections

Lesion-induced direct structural disconnections should lead to increased SSPLs relative to the normative values derived from the HCP tractography atlas (**Fig. 1b**), and accordingly, patients showed reductions in short (i.e. < 3) SSPLs and increases in long (i.e. > 4) SSPLs (**Fig. S1a**). Across patients, the total number of indirect structural disconnections was strongly related to the total number of direct structural disconnections caused by the lesion (*R*^2^=0.78, *p*<0.001), indicating that patients with more extensive direct structural disconnections also suffered more extensive indirect structural disconnections (**Fig. S1b**).

To characterize the extent of direct and indirect structural disconnections caused by the lesions in our sample, we measured the proportions of directly and indirectly structurally connected region pairs that sustained direct and indirect structural disconnections, respectively. On average across the subset of patients with any direct cortico-cortical structural disconnections (*n*=92), 19.03% of all region pairs with direct cortico-cortical structural connections sustained direct structural disconnections and 20.0% of all region pairs with indirect cortico-cortical structural connections sustained indirect structural disconnections (**Fig. S1c**). These observations indicate that focal brain lesions often have widespread effects on the direct and indirect structural pathways in the brain.

### 3.3 Direct and indirect structural disconnections disrupt functional connectivity

Previous work has shown that normal functional connectivity strengths vary as a function of SSPLs (Goni et al., 2014). Thus, prior to evaluating the effects of direct and indirect structural disconnections on functional connectivity in the patient group, we replicated the previously reported relationship between functional connectivity strengths and SSPLs using the data from our control group. As shown in **Figure S2**, average functional connectivity strengths varied reliably as a function of SSPLs for both interhemispheric and intrahemispheric functional connections that were both within and between resting-state networks, and this effect was independent of inter-regional distances.

Next, we proceeded to assess and compare the relative effects of direct and indirect structural disconnections on functional connectivity in the patient group. To obtain estimates of functional connectivity disruption at the level of individual connections, we converted each patient’s functional connectivity matrix into a *z*-score matrix using the mean and standard deviation functional connectivity matrices from the control group (**Fig. 3b**). The resulting *z*-score matrices were therefore deviation matrices quantifying the distance (in standard deviation units) of each functional connection from the expected value in healthy controls (**Fig. 3b**). For each patient, functional connectivity *z*-scores were separately averaged across the patient-specific sets of regions with spared direct structural connections, direct structural disconnections, spared indirect structural connections, and indirect structural connections (see **Fig. 2e**) to obtain a single summary measure of functional connectivity disruption for each set of patient-specific regions.

We performed a three-way repeated-measures analysis of variance (ANOVA) to compare mean functional connectivity *z*-scores among sets of regions with different structural connection types (i.e. direct vs. indirect), structural connection statuses (i.e. spared vs. disconnected), and normative functional connectivity signs as indicated by the mean control functional connectivity matrix (i.e. positive vs. negative; see **Fig. 3b; Fig. 4a,c**) in the patient group. Normative functional connectivity signs were included as a factor because prior work has shown that lesion-induced disruptions of positive and negative functional connections tend to be in opposite directions (Baldassarre et al., 2014; Griffis et al., 2019; Lim et al., 2014; Siegel et al., 2016), and this allowed us to avoid mixing effects that we expected *a priori* to differ in directionality. Within-network functional connections overwhelmingly had positive signs, while the signs of between-network functional connections varied depending on the specific networks in question (**Fig. 4a,c**).

**Fig. 4.**
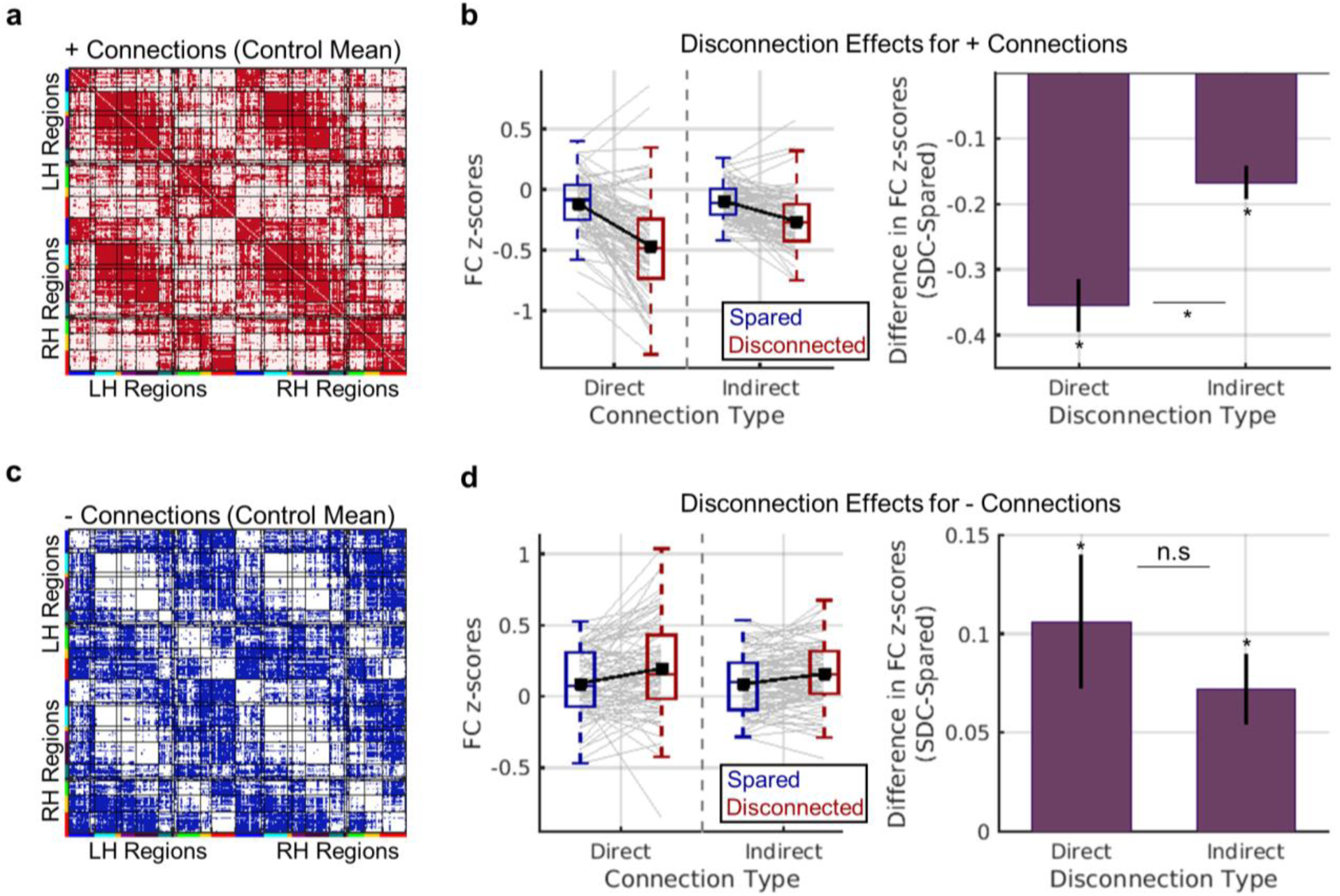
Effects of direct and indirect disconnection on functional connectivity. **(a-b)** Results for positive functional connections. **(a)** The matrix shows positive functional connections from the mean control functional connectivity matrix in red. The analyses shown in (b) were restricted to these connections. **(b)** Left – Distributions of mean patient-level functional connectivity *z*-scores (y-axis) for each connection status (blue vs. red boxplots) and connection type (x-axis). Grey line plots correspond to individual patient observations, and black line plots show group-level means. Right – Data are summarized to show group-level mean (+/-SEM) differences in functional connectivity *z*-scores (y-axis) between regions with disconnected vs. spared direct and indirect connections (x-axis). **(c-d)** Results for negative functional connections. **(c)** The matrix shows negative functional connections from the mean control functional connectivity matrix in blue. The analyses shown in (d) were restricted to these connections. **(d)** Same as (b), but for negative functional connections. *dependent samples *t*-test FWE*p*<0.004. See also **Figs. S3-S4.**

The three-way ANOVA, which included 84 patients with valid data for all factor combinations, revealed a significant three-way interaction of connection status, connection type, and normative FC sign (Greenhouse-Geisser corrected *F*_.34,27.54_=28.09, *p*<0.001). We performed follow-up dependent samples *t-*tests with Bonferroni-Holm correction to further interrogate the effects of interest while controlling the family-wise error rate at 0.05 (Aickin and Gensler, 1996). For positive functional connections (**Fig. 4a**), mean functional connectivity *z*-scores were significantly more negative for regions that sustained structural disconnections than for regions with spared structural connections (**Fig. 4b**). While this effect was observed for both direct and indirect structural disconnection types, the effect was greater for direct structural disconnections than for indirect structural disconnections (**Fig. 4b**, right plot). For negative functional connections (**Fig. 4c**), mean functional connectivity *z*-scores were significantly more positive for regions that sustained structural disconnections than for regions with spared sustained connections (**Fig. 4d**). While this effect was again observed for both direct and indirect structural disconnection types, the magnitudes of direct and indirect structural disconnection effects on mean functional connectivity *z*-scores did not significantly differ for negative functional connections (**Fig. 4d**, right plot). These results support the conclusion that lesion-induced structural disconnections of intermediary structural connections represent a general mechanism underlying lesion-induced functional connectivity disruptions between indirectly structurally connected region pairs across the brain.

Consistent effects of direct and indirect structural disconnections on functional connectivity were also observed when data were further summarized by network (within vs. between) and hemispheric (interhemispheric vs. intrahemispheric) connection types (**Fig. S3**). Supplemental analyses further indicated that the observed results could not be attributed to regional GM damage, vascular/hemodynamic abnormalities as reflected by large hemodynamic lags (Siegel et al., 2017, 2015), or the use of healthy controls as a reference population (**Fig. S4**).

## 4. Discussion

It has long been recognized that focal brain lesions have distributed functional consequences (Carrera and Tononi, 2014). Over the past decade, a number of studies have documented lesion-induced disruptions of resting-state functional connectivity (Eldaief et al., 2016; Gratton et al., 2012; Nomura et al., 2010; Ovadia-Caro et al., 2013; Yuan et al., 2017) and linked them to behavioral deficits in clinical populations (Carter et al., 2010; He et al., 2007; Park et al., 2011; Siegel et al., 2016; Tang et al., 2016). Here, we aimed to clarify how these widespread disruptions of functional connectivity relate to the effects of the lesion on the underlying structural connectome.

Our results show that focal brain lesions have wide-reaching effects on both direct and indirect structural connections, such that roughly one-fifth of all region pairs in the brain suffered either direct or indirect structural connections on average in our patient sample. Further, they show that these lesion-induced changes in the structural connectome are biologically relevant, as they are reflected by disruptions of resting-state functional connectivity between both directly and indirectly disconnected regions. Altogether, the current results indicate that lesion-induced structural disconnections exert a combination of direct and indirect effects on the structural connectome that disrupt ongoing inter-regional signaling and manifest as disruptions of resting-state functional connectivity.

### 4.1. Functional connectivity depends on direct and indirect structural connections

Previous work has shown that normal functional connectivity depends on both the direct and indirect structural connections between brain regions (Adachi et al., 2011; Goni et al., 2014; Honey and Sporns, 2009; Mišić et al., 2015). In the healthy brain, it has been observed that functional connectivity tends to be strongest between regions with direct connections (i.e. SSPLs=1) and weakest between regions with long SSPLs (Goni et al., 2014), although other factors such as the network embedding of structural paths likely also play important roles in determining normative functional connectivity (Goni et al., 2014; Osmanlioğlu et al., 2019). Importantly, we were able to replicate previously reported SSPL effects on functional connectivity in our healthy control data (**Fig. S2**), providing additional support for the conclusion that structural connections, both direct and indirect, shape normal functional connectivity patterns.

### 4.2. Disruptions of the structural connectome are reflected in the functional connectome

Despite the clear dependence of functional connectivity on structural connectivity in the healthy brain, most prior empirical work relating lesion-induced disruptions of functional connectivity to the underlying structural damage has focused on the effects of damage to grey matter regions with particular functional network affiliations (Eldaief et al., 2016; Nomura et al., 2010; Ovadia-Caro et al., 2013) or hub-like functional connectivity profiles (Gratton et al., 2012). In contrast, we recently reported that information about direct structural disconnections is superior to information about lesion size, lesion location, or damage to putative hub regions for explaining core network-level functional connectivity disruptions associated with stroke (Griffis et al., 2019). The results reported here extend our previous work by showing that direct structural connections reliably disrupt resting-state functional connectivity between the disconnected region pairs (**Fig. 4**, see “direct”). Importantly, they also extend previous work on the up/downstream consequences of focal brain lesions (Lu et al., 2011) by showing that damage to intermediary structural connections along the shortest path linking indirectly structurally connected region pairs represents a general mechanism underlying lesion-induced functional connectivity disruptions across the cortex (**Fig. 4**, see “indirect”).

Direct structural disconnections tended to produce more severe functional connectivity disruptions than indirect structural disconnections for positive functional connections (**Fig. 4a**). The results of our supplemental analyses suggest that this effect was likely driven primarily by interhemispheric disconnections (**Fig. S3b**), which are known to rely heavily on direct anatomical projections (Johnston et al., 2008; Roland et al., 2017; Shen et al., 2015). This suggests that the relative magnitudes of functional connectivity disruptions that result from direct and indirect disconnections likely depend on the nature of the affected connections. Determining what factors influence the relative vulnerabilities of different types of functional connections to direct and indirect disconnections is therefore an important goal for future studies on this topic.

Recent studies have shown that lesion-induced direct structural disconnections are associated with neurodegenerative changes in directly disconnected cortical areas (Duering et al., 2015; Foulon et al., 2018; Kuceyeski et al., 2014). For example, cortical thinning has been reported to occur in directly disconnected areas over the first 6 months of recovery (Duering et al., 2015). Speculatively, these regional structural changes might be long-term consequences of disconnection-induced functional disruptions such as those reported here, and this should be addressed by future work. Given that we also observed clear effects of indirect disconnection on functional connectivity, future studies on how structural disconnections impact local regional structure might also incorporate measures of indirect structural disconnection to assess the relative contributions of direct and indirect structural disconnections to changes in local regional structure associated with focal brain lesions.

### 4.3. Potential clinical implications

Our results have potential implications for the development and refinement of techniques that aim to restore brain network function after stroke. For example, there is growing interest in applying non-invasive neuromodulatory techniques such as transcranial magnetic stimulation (TMS) to facilitate stroke recovery (Grefkes and Ward, 2014; Hesse et al., 2011; Naeser et al., 2012), and it has been previously proposed that the putative benefits of techniques such as TMS for stroke recovery may be dependent on the post-stroke structural network architecture (Grefkes and Ward, 2014). The results of our analyses indicate that patient-specific patterns of direct and indirect structural disconnections provide relevant information about the post-stroke structural and functional connectomes. This type of information could potentially be used to both (1) identify potential TMS targets based on their residual post-stroke connectivity patterns and (2) determine how the effects of TMS depend on the attributes of the post-stroke structural connectome.

However, a primary limitation of the current work is that the inferences are necessarily restricted to the group-level rather than at the level of individual patients. That is, while we show that sets of disconnected regions on average show more severe functional connectivity disruptions than sets of regions with spared connections across patients, we do not show that our disconnection measures are sufficient to allow for the accurate prediction of connection-level functional connectivity disruptions in an individual patient, which will likely be necessary to enable the translation of these findings to clinical applications. Dedicated computational modeling (Adhikari et al., 2017) and/or predictive modeling (Goni et al., 2014) studies that incorporate comprehensive disconnection information are therefore an important next step, as are studies that compare the relative utility of atlas-based vs. patient-based measures of disconnection.

Further, while changes in SSPLs computed based on binary structural connectivity matrices, such as those measured here, provide a simple and straightforward means of estimating the indirect consequences of focal brain lesions, they may be too coarse of a measure to allow for direct practical application. More complex network measures such as search information (Goni et al., 2014) or communicability (Osmanlioğlu et al., 2019) may provide more accurate descriptions of the higher-order structural network topology, which may be necessary to enable clinical translation. Ultimately, further research is necessary to further develop and refine these measures, and dedicated experimental studies are necessary to determine the practical utility of this type of information in the context of non-invasive neuromodulation and other potential clinical applications.

## Acknowledgments

Data were provided [in part] by the Human Connectome Project, WU-Minn Consortium (Principal Investigators: David Van Essen and Kamil Ugurbil; 1U54MH091657) funded by the 16 NIH Institutes and Centers that support the NIH Blueprint for Neuroscience Research; and by the McDonnell Center for Systems Neuroscience at Washington University. We thank Alexandre Carter for assisting with lesion segmentation.

## Author Contributions

J.G and G.S designed the analyses and wrote the paper. J.G and N.M. performed data processing and analyses. J.G., G.S., and M.C. edited the paper. G.S. and M.C. contributed data and other resources.

## Supplemental Material

**Fig. S1.**
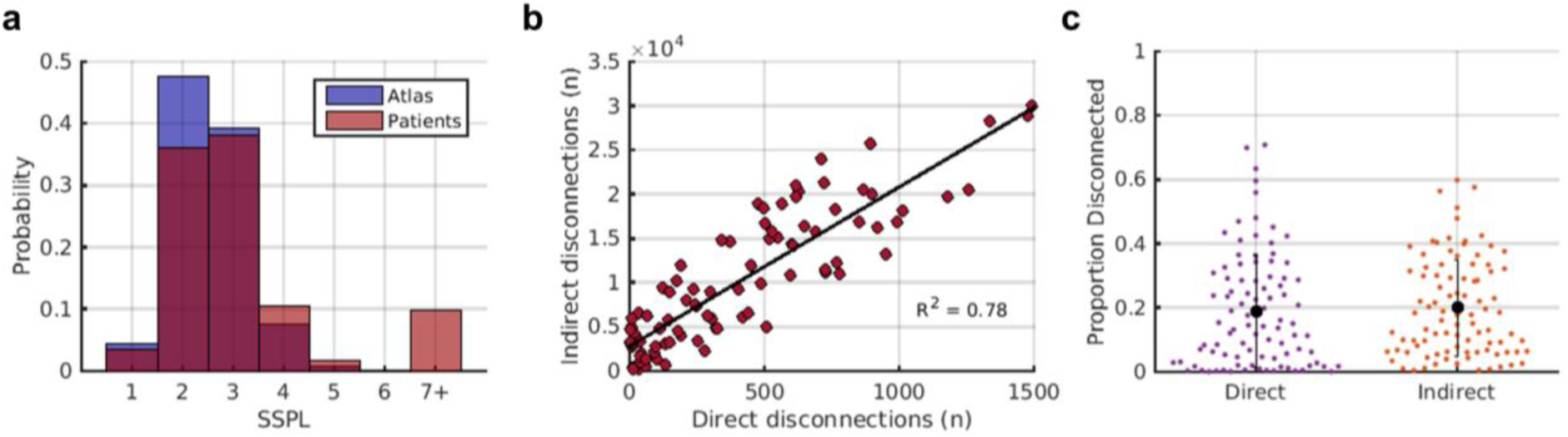
Effects of lesions on SSPLs and relationship between extent of direct and indirect disconnections. **(a)** The red histogram shows the distribution of SSPLs in the atlas-derived structural connectome. The blue histogram shows the distribution of SSPLs in patients. Short SSPLs are decreased and long SSPLs are increased in patients relative to the atlas-derived structural connectome. **(b)** The scatterplot shows the relationship between the total number of direct disconnections (x-axis) and the total number of indirect disconnections (y-axis) in patients. **(c)** Distributions and means/standard deviations are shown for the proportion of disconnected direct and indirect cortico-cortical structural connections in the subset of patients that sustained at least one direct cortico-cortical disconnection (*n*=92). Dots correspond to individual patients.

**Fig. S2.**
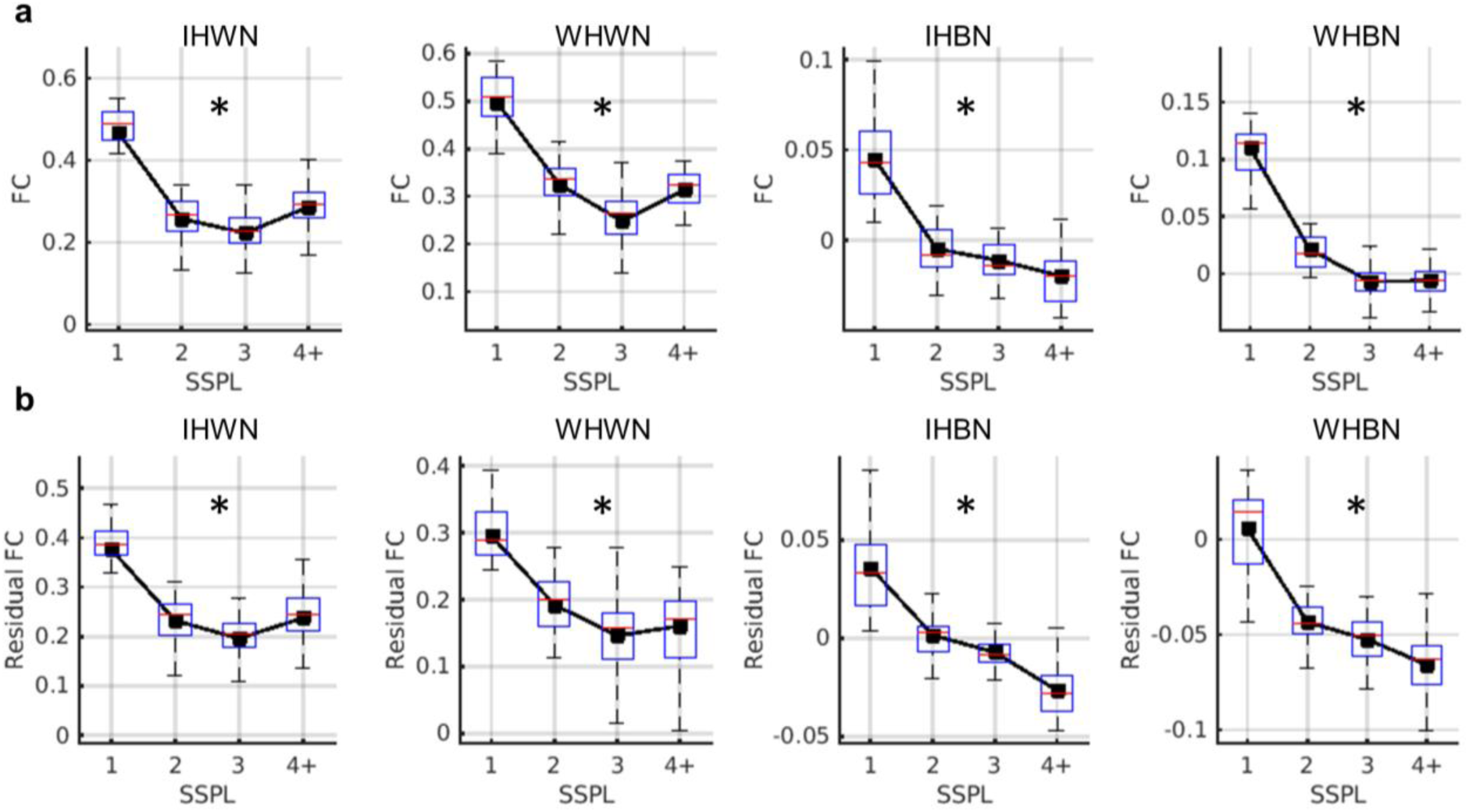
SSPL effects on normal functional connectivity for different network and hemispheric connection types. **(a-b).** Distributions of mean functional connectivity values (y-axis) for region pairs with different atlas-derived SSPLs (x-axis) in healthy controls (box plots) for interhemispheric within-network (IHWN), intrahemispheric within-network (IHWN), interhemispheric between-network (IHBN), and intrahemispheric between-network (WHBN) connection types. The plot in (a) shows the effect without distance regression, while the plots in (b) show the effects after regressing out the effects of spatial distance from each participants functional connectivity matrix. *Greenhouse-Geisser corrected repeated measures ANOVA with factor of SSPL, p<0.001

**Fig. S3.**
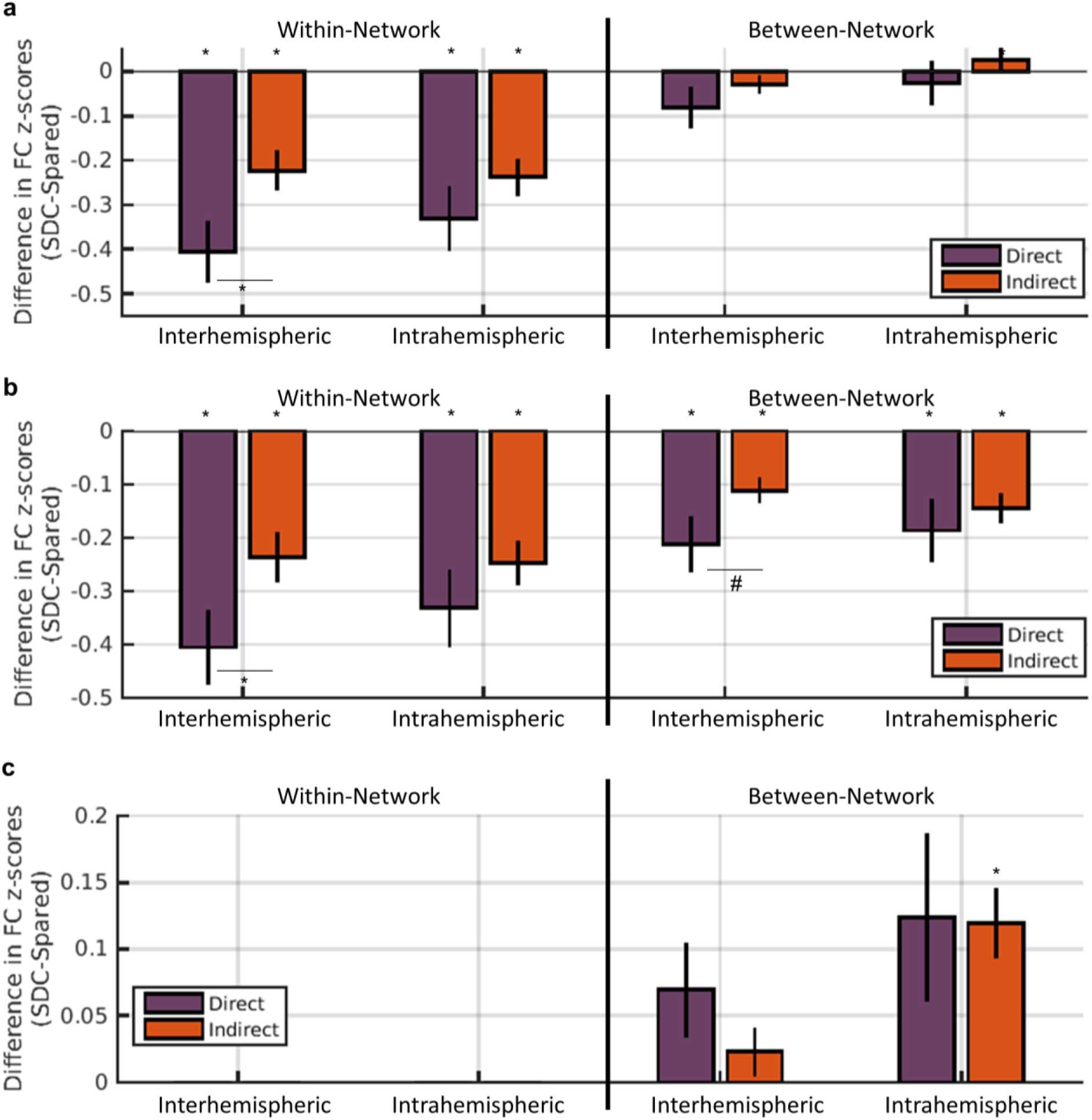
Network and hemispheric connectivity type analyses. **(a-c)** Bar plots show mean (+/-SEM) differences in functional connectivity z-scores (y-axis) between disconnected and spared regions with direct (purple) and indirect (orange) structural connections for each network connection type (x-axis). Plots show analyses using all functional connections (a), only connections with positive normative functional connectivity signs (b), and only connections with negative normative functional connectivity signs (c). Note, for analyses in (c), there were insufficient observations for within-network connections. *dependent samples t-test FWEp<0.05, #dependent samples t-test FWEp=0.09.

**Fig. S4.**
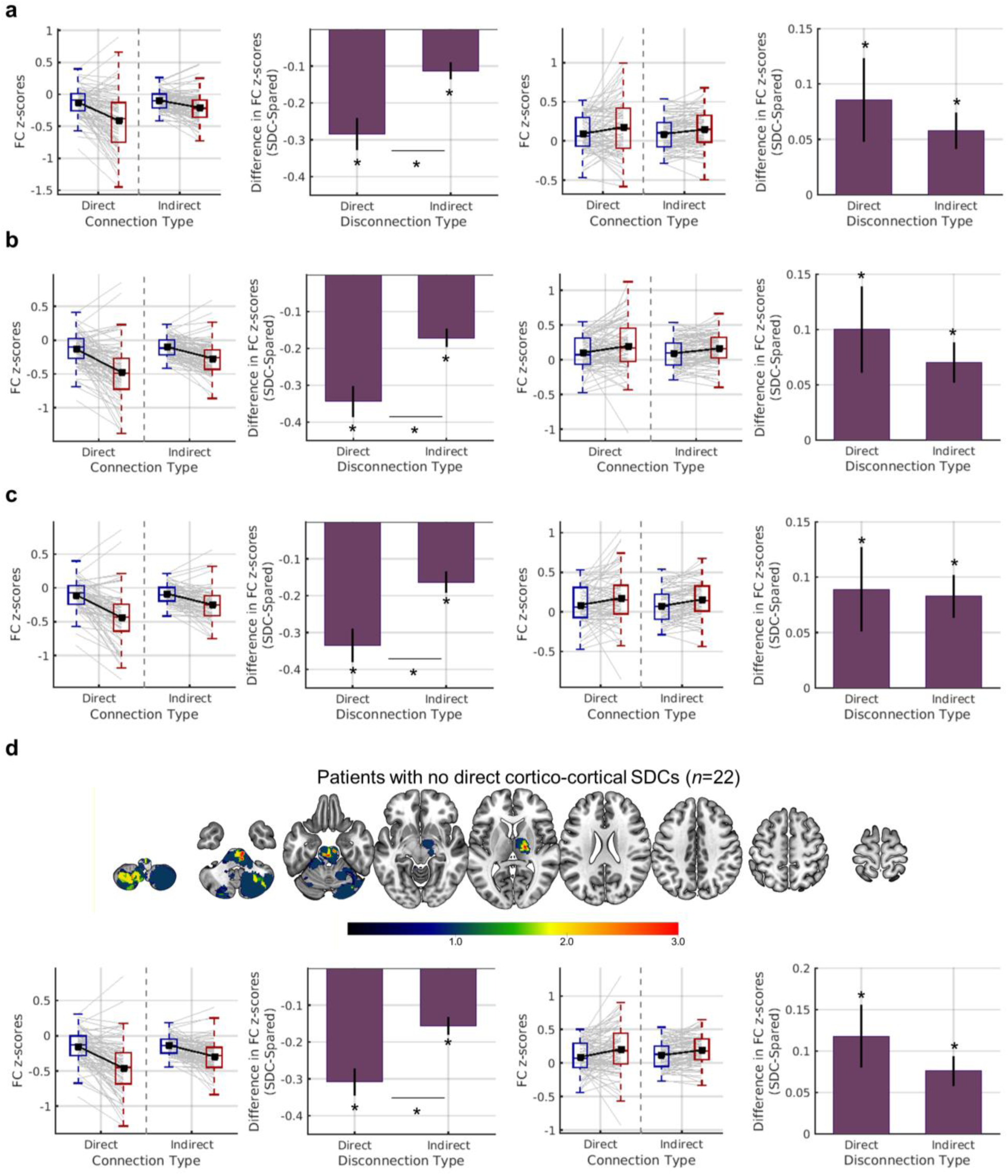
Control analyses. **(a-d).** The plots show the effects obtained when analyses were performed (a) after excluding any damaged regions, (b) using a 50% disconnection threshold for determining direct disconnection, (c) after excluding patients with mean interhemispheric hemodynamic lag differences > 2 standard deviations above the mean observed in controls, and (d) when a sample of 22 stroke patients with no direct cortico-cortical disconnections were used as the reference group for computing functional connectivity z-scores rather than the healthy control group. The lesion overlay for the group of patients with no direct cortico-cortical disconnections is shown in the top panel of (d). Plots on the left show results for positive functional connections, and plots on the right show results for negative functional connections.

## References

Abdi, H., 2010. The greenhouse-geisser correction. Encycl. Res. Des. Sage Publ. 1–10. https://doi.org/10.1007/BF02289823

Adachi, Y., Osada, T., Sporns, O., 2011. Functional connectivity between anatomically unconnected areas is shaped by collective network-level effects in the macaque cortex. Cereb. cortex 1586–1592. https://doi.org/10.1093/cercor/bhr234

Adhikari, M.H., Hacker, C.D., Siegel, J.S., Griffa, A., Hagmann, P., Deco, G., Corbetta, M., 2017. Decreased integration and information capacity in stroke measured by whole brain models of resting state activity. Brain 1068–1085. https://doi.org/10.1093/brain/awx021

Aickin, M., Gensler, H., 1996. Adjusting for multiple testing when reporting research results: The Bonferroni vs Holm methods. Am. J. Public Health 86, 726–728. https://doi.org/10.2105/AJPH.86.5.726

Baldassarre, A., Ramsey, L., Hacker, C.L., Callejas, A., Astafiev, S. V., Metcalf, N. V., Zinn, K., Rengachary, J., Snyder, A.Z., Carter, A.R., Shulman, G.L., Corbetta, M., 2014. Large-scale changes in network interactions as a physiological signature of spatial neglect. Brain 137, 3267–3283. https://doi.org/10.1093/brain/awu297

Baldassarre, A., Ramsey, L.E., Siegel, J.S., 2016. Brain connectivity and neurological disorders after stroke. Curr. Opin. Neurol. 29, 706–713. https://doi.org/10.1097/WCO.0000000000000396

Biswal, B., Zerrin, Y., Haughton, V.M., Hyde, J.S., 1995. Functional Connectivity in the Motor Cortex of Resting Human Brain Using Echo-Planar MRI. Magn. Reson. Med. 34. https://doi.org/10.1002/mrm.1910340409

Carrera, E., Tononi, G., 2014. Diaschisis: past, present, future. Brain 137, 2408–2422. https://doi.org/10.1093/brain/awu101

Carter, A.R., Astafiev, S. V., Lang, C.E., Connor, L.T., Rengachary, J., Strube, M.J., Pope, D.L.W., Shulman, G.L., Corbetta, M., 2010. Resting interhemispheric functional magnetic resonance imaging connectivity predicts performance after stroke. Ann. Neurol. 67, 365–375. https://doi.org/10.1002/ana.21905

Corbetta, M., Ramsey, L., Callejas, A., Baldassarre, A., Hacker, C.D.D., Siegel, J.S.S., Astafiev, S.V. V., Rengachary, J., Zinn, K., Lang, C.E.E., Connor, L.T.T., Fucetola, R., Strube, M., Carter, A.R.R., Shulman, G.L.L., 2015. Common Behavioral Clusters and Subcortical Anatomy in Stroke. Neuron 85, 927–941. https://doi.org/10.1016/j.neuron.2015.02.027

Dale, A.M., Fischl, B., Sereno, M.I., 1999. Cortical surface-based analysis. I. Segmentation and surface reconstruction. Neuroimage 9, 179–94. https://doi.org/10.1006/nimg.1998.0395

Duering, M., Righart, R., Wollenweber, F.A., Zietemann, V., Gesierich, B., Dichgans, M., 2015. Acute infarcts cause focal thinning in remote cortex via degeneration of connecting fiber tracts. Neurology 84, 1685–1692. https://doi.org/10.1212/WNL.0000000000001502

Eldaief, M.C., McMains, S., Hutchison, R.M., Halko, M.A., Pascual-Leone, A., 2016. Reconfiguration of Intrinsic Functional Coupling Patterns Following Circumscribed Network Lesions. Cereb. Cortex bhw139. https://doi.org/10.1093/cercor/bhw139

Fischl, B., Sereno, M.I., Dale, A.M., 1999. Cortical surface-based analysis. II: Inflation, flattening, and a surface-based coordinate system. Neuroimage 9, 195–207. https://doi.org/10.1006/nimg.1998.0396

Foulon, C., Cerliani, L., Kinkingnéhun, S., Levy, R., Rosso, C., Urbanski, M., Volle, E., Thiebaut de Schotten, M., Kinkingnehun, S., Levy, R., Rosso, C., Urbanski, M., Volle, E., Thiebaut de Schotten, M., Kinkingnéhun, S., Levy, R., Rosso, C., Urbanski, M., Volle, E., de Schotten, M.T., 2018. Advanced Lesion Symptom Mapping Analyses And Implementation As BCBtoolkit. Gigascience 7, 1–17. https://doi.org/10.1101/133314

Glasser, M.F., Sotiropoulos, S.N., Wilson, J.A., Coalson, T.S., Fischl, B., Andersson, J.L., Xu, J., Jbabdi, S., Webster, M., Polimeni, J.R., Van Essen, D.C., Jenkinson, M., 2013. The minimal preprocessing pipelines for the Human Connectome Project. Neuroimage 80, 105–124. https://doi.org/10.1016/j.neuroimage.2013.04.127

Goni, J., van den Heuvel, M.P., Avena-Koenigsberger, A., Velez de Mendizabal, N., Betzel, R.F., Griffa, A., Hagmann, P., Corominas-Murtra, B., Thiran, J.-P., Sporns, O., 2014. Resting-brain functional connectivity predicted by analytic measures of network communication. Proc. Natl. Acad. Sci. 111, 833–838. https://doi.org/10.1073/pnas.1315529111

Gordon, E.M., Laumann, T.O., Adeyemo, B., Huckins, J.F., Kelley, W.M., Petersen, S.E., 2016. Generation and Evaluation of a Cortical Area Parcellation from Resting-State Correlations. Cereb. Cortex 26, 288–303. https://doi.org/10.1093/cercor/bhu239

Gratton, C., Nomura, E.M., Pérez, F., D’Esposito, M., 2012. Focal brain lesions to critical locations cause widespread disruption of the modular organization of the brain. J. Cogn. Neurosci. 24, 1275–1285. https://doi.org/10.1162/jocn_a_00222

Grefkes, C., Ward, N.S., 2014. Cortical reorganization after stroke: how much and how functional? Neuroscientist 20, 56–70. https://doi.org/10.1177/1073858413491147

Griffis, J.C., Metcalf, N. V, Corbetta, M., Shulman, G.L., 2019. Structural disconnections explain brain network dysfunction after stroke. Cell Rep. 28, 1–68. https://doi.org/10.1101/562165

He, B.J., Snyder, A.Z., Vincent, J.L., Epstein, A., Shulman, G.L., Corbetta, M., 2007. Breakdown of functional connectivity in frontoparietal networks underlies behavioral deficits in spatial neglect. Neuron 53, 905–18. https://doi.org/10.1016/j.neuron.2007.02.013

Hesse, M.D., Sparing, R., Fink, G.R., 2011. Ameliorating spatial neglect with non-invasive brain stimulation: From pathophysiological concepts to novel treatment strategies. Neuropsychol. Rehabil. 21, 676–702. https://doi.org/10.1080/09602011.2011.573931

Honey, C., Sporns, O., 2009. Predicting human resting-state functional connectivity from structural connectivity. Proc. … 106, 1–6.

Johnston, J.M., Vaishnavi, S.N., Smyth, M.D., Zhang, D., He, B.J., Zempel, J.M., Shimony, J.S., Snyder, A.Z., Raichle, M.E., 2008. Loss of Resting Interhemispheric Functional Connectivity after Complete Section of the Corpus Callosum. J. Neurosci. 28, 6453–6458. https://doi.org/10.1523/JNEUROSCI.0573-08.2008

Kuceyeski, A., Kamel, H., Navi, B.B., Raj, A., Iadecola, C., 2014. Predicting future brain tissue loss from white matter connectivity disruption in ischemic stroke. Stroke 45, 717–722. https://doi.org/10.1161/STROKEAHA.113.003645

Lim, D.H., LeDue, J.M., Mohajerani, M.H., Murphy, T.H., 2014. Optogenetic Mapping after Stroke Reveals Network-Wide Scaling of Functional Connections and Heterogeneous Recovery of the Peri-Infarct. J. Neurosci. 34, 16455–16466. https://doi.org/10.1523/JNEUROSCI.3384-14.2014

Lu, J., Liu, H., Zhang, M., Wang, D., Cao, Y., Ma, Q., Rong, D., Wang, X., Buckner, R.L., Li, K., 2011. Focal Pontine Lesions Provide Evidence That Intrinsic Functional Connectivity Reflects Polysynaptic Anatomical Pathways. J. Neurosci. 31, 15065–15071. https://doi.org/10.1523/JNEUROSCI.2364-11.2011

Mišić, B., Betzel, R.F., Nematzadeh, A., Goñi, J., Griffa, A., Hagmann, P., Flammini, A., Ahn, Y.Y., Sporns, O., 2015. Cooperative and Competitive Spreading Dynamics on the Human Connectome. Neuron 86, 1518–1529. https://doi.org/10.1016/j.neuron.2015.05.035

Naeser, M. a, Martin, P.I., Ho, M., Treglia, E., Kaplan, E., Bashir, S., Pascual-Leone, A., 2012. Transcranial magnetic stimulation and aphasia rehabilitation. Arch. Phys. Med. Rehabil. 93, S26–34. https://doi.org/10.1016/j.apmr.2011.04.026

Nomura, E.M., Gratton, C., Visser, R.M., Kayser, A., Perez, F., D’Esposito, M., 2010. Double dissociation of two cognitive control networks in patients with focal brain lesions. Proc. Natl. Acad. Sci. 107, 12017–12022. https://doi.org/10.1073/pnas.1002431107

Osmanlioğlu, Y., Tunç, B., Parker, D., Elliott, M.A., Baum, G.L., Ciric, R., Satterthwaite, T.D., Gur, R.E., Gur, R.C., Verma, R., 2019. System-level matching of structural and functional connectomes in the human brain. Neuroimage 199, 93–104. https://doi.org/10.1016/j.neuroimage.2019.05.064

Ovadia-Caro, S., Villringer, K., Fiebach, J., Jungehulsing, G.J., van der Meer, E., Margulies, D.S., Villringer, A., 2013. Longitudinal effects of lesions on functional networks after stroke. J. Cereb. Blood Flow Metab. 33, 1279–85. https://doi.org/10.1038/jcbfm.2013.80

Park, C., Chang, W.H., Ohn, S.H., Kim, S.T., Bang, O.Y., Pascual-Leone, A., Kim, Y.-H., 2011. Longitudinal changes of resting-state functional connectivity during motor recovery after stroke. Stroke. 42, 1357–62. https://doi.org/10.1161/STROKEAHA.110.596155

Power, J., Mitra, A., Laumann, T.O., Snyder, A.Z., Schlaggar, B.L., Petersen, S.E., 2014. Methods to detect, characterize, and remove motion artifact in resting state fMRI. Neuroimage 84, 320–341. https://doi.org/10.1016/j.neuroimage.2013.08.048

Pustina, D., Avants, B., Faseyitan, O., Medaglia, J., Branch Coslett, H., 2017. Improved accuracy of lesion to symptom mapping with multivariate sparse canonical correlations. Neuropsychologia 8000, 1–13. https://doi.org/10.1016/j.neuropsychologia.2017.08.027

Raffin, E., Siebner, H.R., 2014. Transcranial brain stimulation to promote functional recovery after stroke. Curr. Opin. Neurol. 27, 54–60. https://doi.org/10.1097/WCO.0000000000000059

Robb, R.A., Hanson, D.P., 1991. A software system for interactive and quantitative visualization of multidimensional biomedical images. Australas. Phys. Eng. Sci. Med.

Roland, J.L., Snyder, A.Z., Hacker, C.D., Mitra, A., Shimony, J.S., Limbrick, D.D., Raichle, M.E., Smyth, M.D., Leuthardt, E.C., 2017. On the role of the corpus callosum in interhemispheric functional connectivity in humans. Proc. Natl. Acad. Sci. 114, 201707050. https://doi.org/10.1073/pnas.1707050114

Rubinov, M., Sporns, O., 2010. Complex network measures of brain connectivity: uses and interpretations. Neuroimage 52, 1059–69. https://doi.org/10.1016/j.neuroimage.2009.10.003

Shen, K., Mišić, B., Cipollini, B.N., Bezgin, G., Buschkuehl, M., Hutchison, R.M., Jaeggi, S.M., Kross, E., Peltier, S.J., Everling, S., Jonides, J., McIntosh, A.R., Berman, M.G., 2015. Stable long-range interhemispheric coordination is supported by direct anatomical projections. Proc. Natl. Acad. Sci. 112, 6473–6478. https://doi.org/10.1073/pnas.1503436112

Siegel, J.S., Ramsey, L.E., Snyder, A.Z., Metcalf, N. V, Chacko, R. V, Weinberger, K., Baldassarre, A., Hacker, C., Shulman, G.L., Corbetta, M., 2016. Disruptions of network connectivity predict impairment in multiple behavioral domains after stroke. PNAS I, 1–10. https://doi.org/10.1073/pnas.1521083113

Siegel, J.S., Seitzman, B.A., Ramsey, L.E., Ortega, M., Gordon, E.M., Dosenbach, N.U.F., Petersen, S.E., Shulman, G.L., Corbetta, M., 2018. Re-emergence of modular brain networks in stroke recovery. Cortex 101, 44–59. https://doi.org/10.1016/j.cortex.2017.12.019

Siegel, J.S., Shulman, G.L., Corbetta, M., 2017. Measuring functional connectivity in stroke: Approaches and considerations. J. Cereb. Blood Flow Metab. 0271678X1770919. https://doi.org/10.1177/0271678X17709198

Siegel, J.S., Snyder, A.Z., Ramsey, L., Shulman, G.L., Corbetta, M., 2015. The effects of hemodynamic lag on functional connectivity and behavior after stroke. J. Cereb. Blood Flow Metab. 0271678X15614846. https://doi.org/10.1177/0271678X15614846

Tang, C., Zhao, Z., Chen, C., Zheng, X., Sun, F., Zhang, X., Tian, J., Fan, M., Wu, Y., Jia, J., 2016. Decreased Functional Connectivity of Homotopic Brain Regions in Chronic Stroke Patients: A Resting State fMRI Study. PLoS One 11, 1–13. https://doi.org/10.1371/journal.pone.0152875

Tzourio-Mazoyer, N., Landeau, B., Papathanassiou, D., Crivello, F., Etard, O., Delcroix, N., Mazoyer, B., Joliot, M., 2002. Automated anatomical labeling of activations in SPM using a macroscopic anatomical parcellation of the MNI MRI single-subject brain. Neuroimage 15, 273–289. https://doi.org/10.1006/nimg.2001.0978

Van Den Heuvel, M.P., Mandl, R.C.W., Kahn, R.S., Hulshoff Pol, H.E., 2009. Functionally linked resting-state networks reflect the underlying structural connectivity architecture of the human brain. Hum. Brain Mapp. 30, 3127–3141. https://doi.org/10.1002/hbm.20737

Van Essen, D.C., Drury, H.A., Dickson, J., Harwell, J., Hanlon, D., Anderson, C., 2001. An Integrated Software Suite for Surface-based Analyses of Cerebral Cortex. J. Am. Med. Informatics Assoc. 8, 443–459.

Wilson, S.M., Galantucci, S., Tartaglia, M.C., Rising, K., Patterson, D.K., Henry, M.L., Ogar, J.M., DeLeon, J., Miller, B.L., Gorno-Tempini, M.L., 2011. Syntactic processing depends on dorsal language tracts. Neuron 72, 397–403. https://doi.org/10.1016/j.neuron.2011.09.014

Yeh, F.-C., Panesar, S., Fernandes, D., Meola, A., Yoshino, M., Fernandez-Miranda, J.C., Vettel, J.M., Verstynen, T., 2018. Population-averaged atlas of the macroscale human structural connectome and its network topology. Neuroimage 178, 57–68. https://doi.org/10.1016/j.neuroimage.2018.05.027

Yuan, B., Fang, Y., Han, Z., Song, L., He, Y., Bi, Y., 2017. Brain hubs in lesion models: Predicting functional network topology with lesion patterns in patients. Sci. Rep. 7, 17908. https://doi.org/10.1038/s41598-017-17886-x

